# Ketogenesis restrains aging-induced exacerbation of COVID in a mouse model

**DOI:** 10.1101/2020.09.11.294363

**Authors:** Seungjin Ryu, Irina Shchukina, Yun-Hee Youm, Hua Qing, Brandon K. Hilliard, Tamara Dlugos, Xinbo Zhang, Yuki Yasumoto, Carmen J. Booth, Carlos Fernández-Hernando, Yajaira Suárez, Kamal M. Khanna, Tamas L. Horvath, Marcelo O. Dietrich, Maxim N. Artyomov, Andrew Wang, Vishwa Deep Dixit

**Author notes:** Correspondence to (A.W.) or (V.D.D.).

## Abstract

Increasing age is the strongest predictor of risk of COVID-19 severity. Unregulated cytokine storm together with impaired immunometabolic response leads to highest mortality in elderly infected with SARS-CoV-2. To investigate how aging compromises defense against COVID-19, we developed a model of natural murine beta coronavirus (mCoV) infection with mouse hepatitis virus strain MHV-A59 (mCoV-A59) that recapitulated majority of clinical hallmarks of COVID-19. Aged mCoV-A59-infected mice have increased mortality and higher systemic inflammation in the heart, adipose tissue and hypothalamus, including neutrophilia and loss of γδ T cells in lungs. Ketogenic diet increases beta-hydroxybutyrate, expands tissue protective γδ T cells, deactivates the inflammasome and decreases pathogenic monocytes in lungs of infected aged mice. These data underscore the value of mCoV-A59 model to test mechanism and establishes harnessing of the ketogenic immunometabolic checkpoint as a potential treatment against COVID-19 in the elderly.

**Highlights:**

- Natural MHV-A59 mouse coronavirus infection mimics COVID-19 in elderly.
- Aged infected mice have systemic inflammation and inflammasome activation
- Murine beta coronavirus (mCoV) infection results in loss of pulmonary γδ T cells.
- Ketones protect aged mice from infection by reducing inflammation.

**eTOC Blurb:** Elderly have the greatest risk of death from COVID-19. Here, Ryu et al report an aging mouse model of coronavirus infection that recapitulates clinical hallmarks of COVID-19 seen in elderly. The increased severity of infection in aged animals involved increased inflammasome activation and loss of γδ T cells that was corrected by ketogenic diet.

## INTRODUCTION

Aging-driven reduced resilience to infections is dependent in part on the restricted T cell repertoire diversity together with impaired T and B cell activation as well as inflammasome-driven low-grade systemic inflammation that compromises innate immune function (Akbar and Gilroy, 2020; Camell et al., 2017; Youm et al., 2013). Consequently, 80 percent of deaths due to COVID-19 in US are in adults > 65 years old (https://www.cdc.gov/) and aging is the strongest factor to increase infection fatality (Pastor-Barriuso et al., 2020; Perez-Saez et al., 2020; Ward et al., 2020). Lack of an aging animal model that mimics SARS-CoV-2 immunopathology has been a major limitation in the effort to determine the mechanism of disease and to develop effective therapeutics for the elderly. Inability of mouse ACE2 to bind SARS-CoV-2 is a significant hurdle in understanding the basic mechanism of COVID-19. Accordingly, several approaches have been employed to develop models including introduction of human-ACE2 in mice and transient induction of hACE2 through adenoviral-associated vectors. These models have begun to yield important information on the mechanism of disease development. For example, epithelial cell specific induction of hACE2 (K18-hACE2) as a model of SARS-CoV-2 infection demonstrated that post intranasal inoculation, animals develop lung inflammation and pneumonia driven by infiltration of monocytes, neutrophils and T cells (Winkler et al., 2020). Also, initial studies that employ lung ciliated epithelial cell-specific HFH4/FOXJ1 promoter driven hACE2 transgenic mice show SARS-CoV-2 infection induces weight loss, lung inflammation and approximately 50% mortality rate, suggesting the usefulness of this model to understand the mechanism of immune dysregulation (Jiang et al., 2020). However, significant hurdles remain to understand the mechanism and test therapeutic interventions that are relevant to disease severity in elderly, as complicated breeding and specific mutations need to be introduced in hACE2 transgenic strains in addition to the time required to age these models. The mouse model of SARS-CoV-2 based on adeno-associated virus (AAV)–mediated expression of hACE2 may allow circumvention of the above constrains. The delivery of hACE2 into the respiratory tract of C57BL/6 mice with AAV9 causes a productive infection as revealed by >200 fold increase in SARS-CoV-2 RNA and show similar interferon gene expression signatures as COVID-19 patients (Israelow et al., 2020). However, in young wild-type mice, this model induces mild acute respiratory distress syndrome (ARDS) and does not cause neutrophilia, weight loss or lethality (Israelow et al., 2020). Other studies using replication deficient adenovirus-mediated transduction of hACE in mice and infection with SARS-CoV-2 produced 20% weight-loss including lung inflammation (Hassan et al., 2020; Sun et al., 2020). Furthermore, genetic remodeling of the SARS-CoV-2 spike receptor binding domain that allow interaction with mACE demonstrated peribronchiolar lymphocytic inflammatory infiltrates and epithelial damage but no weight-loss in infected mice (Dinnon et al., 2020). Moreover, middle aged female mice (one year old, analogous to approx. 43 year old human), display greater lung pathology and loss of function post infection with 10% weight-loss followed by spontaneous recovery 7 days post infection (Dinnon et al., 2020). However, it remains unclear if this model replicates the clinical, systemic inflammation and immunological response and mortality observed in COVID-19.

The mouse hepatitis virus (MHV) and SARS-CoV-2 are both ARDS-related beta coronaviruses with a high degree of homology (Gorbalenya et al., 2020). Until the emergence of SARS-CoV-2, the natural infection with MHV mouse coronavirus-A59 (mCoV-A59) has traditionally been thought to be of minor relevance to human disease and largely been a primary veterinary concern to maintain the specific-pathogen free status of mouse research facilities (Hickman and Thompson, 2004). However, lack of an aging mouse model of COVID-19 necessitates re-evaluation of mCoV-A59 to investigate the mechanism of multi-organ inflammation, morbidity and mortality caused by the disease. Importantly, the mCoV-A59 utilizes the entry receptor CEACAM1, which is expressed on respiratory epithelium, but also on enterocytes, endothelial cells, and neurons, much like ACE2 (Godfraind et al., 1995) thus allowing the study of wide-ranging systemic impacts of infection. Moreover, natural infection with mCoV-A59 causes ARDS in C57BL/6J animals, while all other MHV strains require the A/J or type-I interferon-deficient background, for the development of severe disease (De Albuquerque et al., 2006; Khanolkar et al., 2009; Yang et al., 2014) limiting their use. Lastly, another major practical advantage of natural infection with mCoV-A59 mouse model is that it does not require limited BSL3 facilities, thus allowing the allocation of precious world-wide BSL3 specialized laboratory space to be prioritized for human SARS-CoV-2 virus studies in primates and other models, which cannot be achieved by using mouse-adapted SARS-CoV-2 models.

Aging-induced chronic inflammation in the absence of overt infections is predominantly driven by the NLRP3 inflammasome (Bauernfeind et al., 2016; Camell et al., 2017; Youm et al., 2013), a myeloid cell-expressed multiprotein complex that senses pathogen associated molecular patterns (PAMPs) and danger associated molecular patterns (DAMPs) to cause the processing and secretion of IL-1β and IL-18. There is increasing evidence that SARS-CoV-2 infection activates the NLRP3 inflammasome (Siu et al., 2019) with increased levels of IL-18 and lactate dehydrogenase (LDH) levels due to inflammasome mediated pyroptotic cell death (Lucas et al., 2020; Zhou et al., 2020). It is now known that increased glycolysis, which activates inflammasome is associated with worsened COVID-19 outcome (Codo et al., 2020). This raises the question whether the substrate switch from glycolysis-to-ketogenesis, can be employed to stave off COVID-19 in high risk elderly population. Here, we establish that intranasal infection with mCoV-A59 recapitulates clinical features of COVID-19 seen in elderly and demonstrate that ketone metabolites protect against disease through inhibition of NLRP3 inflammasome and expansion of protective γδ T cells in lungs.

## RESULTS

### mCoV-A59 infection in aged mice mimics COVID-19 severity

To determine the underlying deficits in immune and inflammatory response in aging, we investigated the impact of mCoV-A59 intranasal inoculation on adult (2-6 month) and old male mice (20-24 month) (Figure 1A). The LD-0 infectious dose of mCoV-A59 in adult (PFU 7e3) caused 100 percent lethality in aged mice (Figure 1B). Compared to adults, the old infected mice displayed greater weight-loss (Figure 1C), hypoxemia (Figure 1D), and anorexia (Figure 1E) without a significant difference in viral load in lungs (Figure S1A). Interestingly, aging led to a significant reduction in CD4, CD4:CD8 ratio (Figure 1F and Figure S1B) and ϒδ T cells (Figure 1G and Figure S1C) in lungs and spleen (Figure S1D) with increased neutrophils (Figure 1H), Ly6C^hi^ monocytes (Figure 1I and S1E) and no change in eosinophils (Figure S1F). In addition, the lungs of old infected mice displayed increased frequency of CD64^+^MerTK^+^ cells (Figure 1J and S1E) with no significant differences in the total population of alveolar or interstitial macrophages (Figure S1G and S1H).

**Figure 1.**
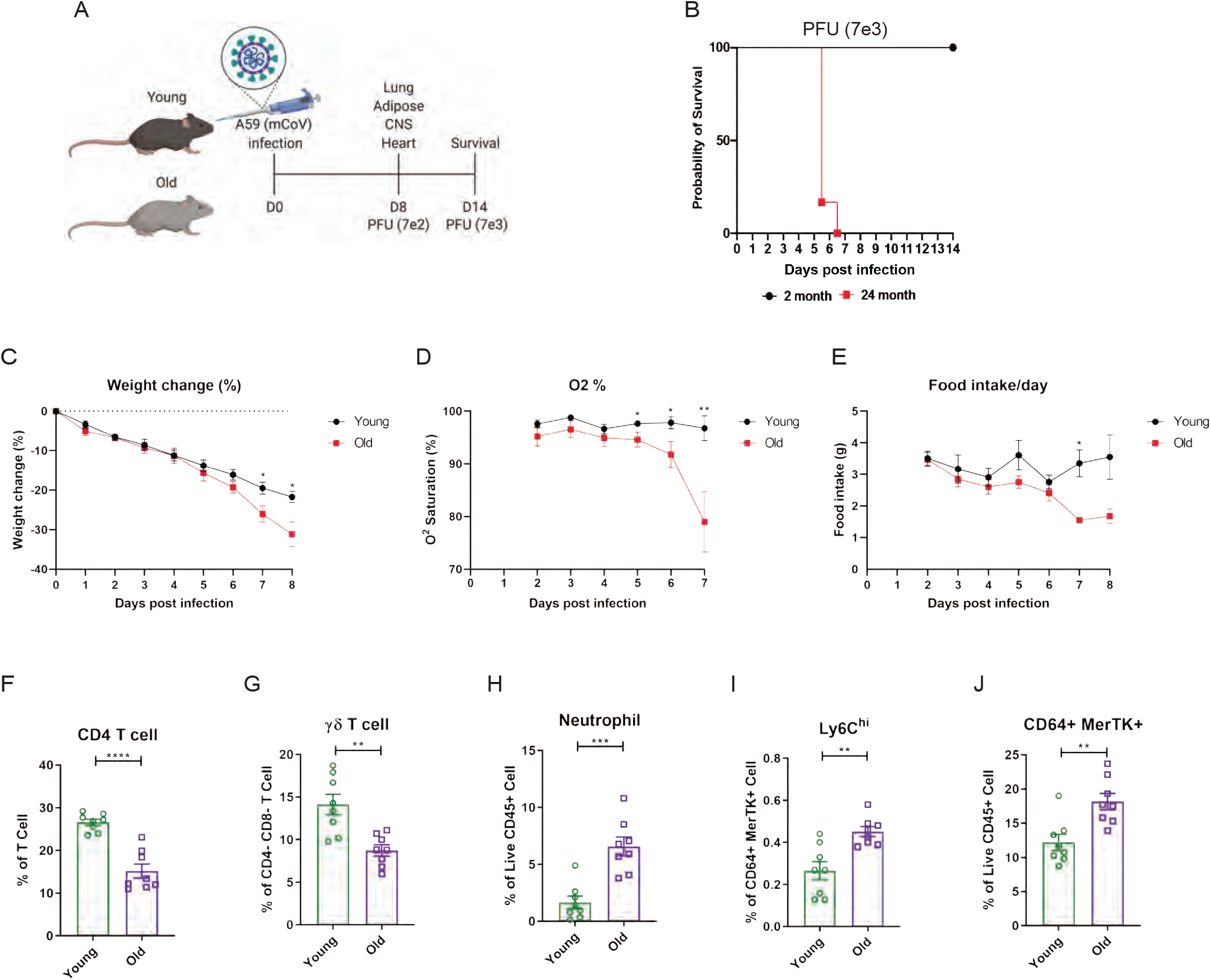
Aging exacerbates A59 (mCoV) infection. **(A)** Schematic of A59 (mCoV) infection experiment with young (2-6 month) and old mice (20-24 month). (**B)** Probability of survival of young (n=6) and old (n=6) infected mice. Survival of mice was examined after infection with high dose of virus (PFU 7e3) up to 14 days. (**C-D)** Young (n=8) and old mice (n=8) were infected with low dose of virus (PFU 7e2), and weight change (%) (C), O_2_ saturation (D), and daily food intake (E) was recorded. (**F-J)** Flow cytometry analysis of of CD4 T cell (F), γδ T cell (G), neutrophil (H), Ly6C^hi^ cell (I), and CD64^+^ MerTK^+^ cell (J) on day 8 post (PFU 7e2) infection. Error bars represent the mean ± S.E.M. Two-tailed unpaired t-tests were performed for statistical analysis. * P < 0.05; ** P < 0.01; *** P < 0.001; **** P < 0.0001.

Transmission electron microscopy confirmed the dissemination of the viral particles in pneumocytes in lungs (Figure 2A). Following mCoV-A59 inoculation, IHC analyses by HE and MSB staining, in both 6 month and 20-24-month old mice, there is perivascular inflammation (arrows, arrowhead) as well as perivascular edema (*) and increased perivascular collagen/fibrosis (MSB, blue) that is more severe in the 20-24-month mCoV-A59 infected mice (Figure 2B). Further, 20-24 month old mice inoculated with mCoV-A59 have dense foci visible at low power (box) and amphophilic material (fibrosis) with few scattered brightly eosinophilic erythrocytes (grey arrowhead) admixed with lymphocytes and plasma cells. By MSB stain, at higher power this same focus (***) in the 20-24 month infected mice revealed that the end of a small blood vessel (BV) terminates in to a mass of collapsed alveoli, without obvious septa admixed with inflammatory cells, disorganized fibrin/collagen fibers (blue) suggestive of ante mortem pulmonary thrombosis in contrast to post mortem blood clots where erythrocytes are yellow (** MSB, yellow) (Figure 2B). Taken together, consistent with ARDS, the lungs of aged mice infected with mCoV-A59 had increased foci of inflammation, immune cell infiltration, perivascular edema, hyaline membrane formation and type II pneumocyte hyperplasia, organizing pneumonia, interstitial pneumonitis, and occasional hemorrhage and microthrombi, affecting approximately 75% of the lungs (Figure 2B).

**Figure 2.**
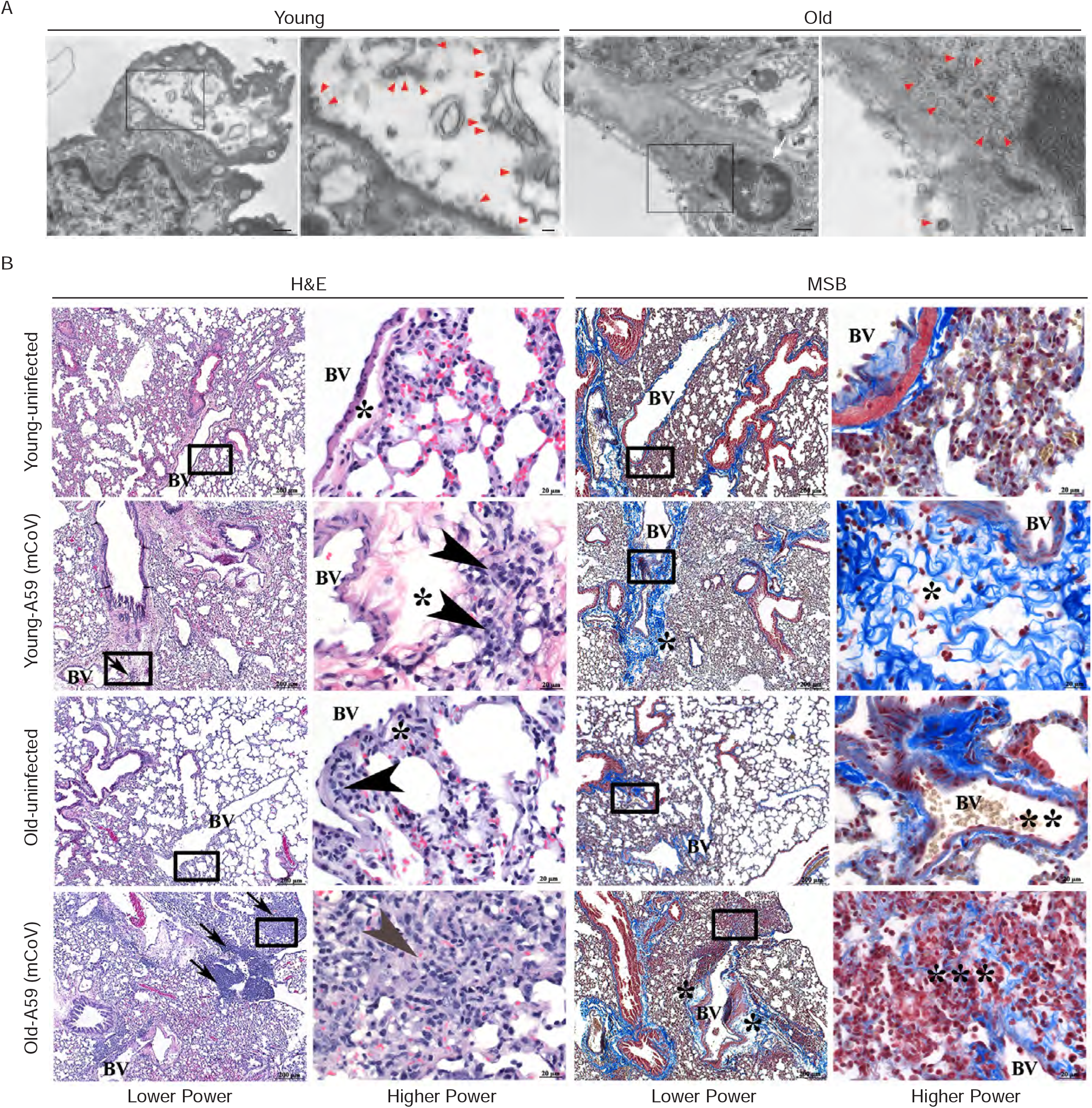
A59 (mCoV) infection significantly affects lung phenotype in old mice. (**A)** Transmission electron microscopic images of A59 (mCoV) particles in pneumocytes. Left panels show pneumocyte of an infected young (top) and old (bottom) mouse. Bar scale represents 500nm. Right panels show zoomed-in images of boxed area of left panels. Pneumocyte with budding viral particles was indicated by red arrowheads. Bar scale represents 100nm. Apoptotic pneumocyte of an infected old mouse showed the shrunken and degraded nucleus (white arrow) and chromatin condensation (white asterisk). (**B)** Representative photomicrographs of hematoxylin and eosin (H&E)-stained (left) and Martius scarlet blue trichrome (MSB)-stained (right) sections of lung from young and old mice 8 days post infection with A59 (mCoV), along with lung from uninfected young and old mice. There are foci of inflammation (arrow), perivascular edema (*), and perivascular lymphocytes and plasma cells (arrow head). Box areas in low power images were used for high power imaging. BV indicates small blood vessel.

### Aging enhances systemic inflammatory response in A59 (mCoV) infected mice

We next investigated whether mCoV-A59 infection in aged mice mimics the hyperinflammatory systemic response seen in elderly patients infected with COVID-19. Compared to young animals, old mice infected with equivalent doses of mCoV-A59 displayed significant increase in circulating IL-1β, TNFα, IL-6 (Figure 3A-C) and MCP-1 without affecting MIP-1β (Figure S2A and S2B). Similar to COVID-19, the infection with mCoV-A59 caused increased cardiac inflammation in old mice as evaluated by greater number of infiltrating CD68^+^ myeloid cells (Figure 3D).

**Figure 3.**
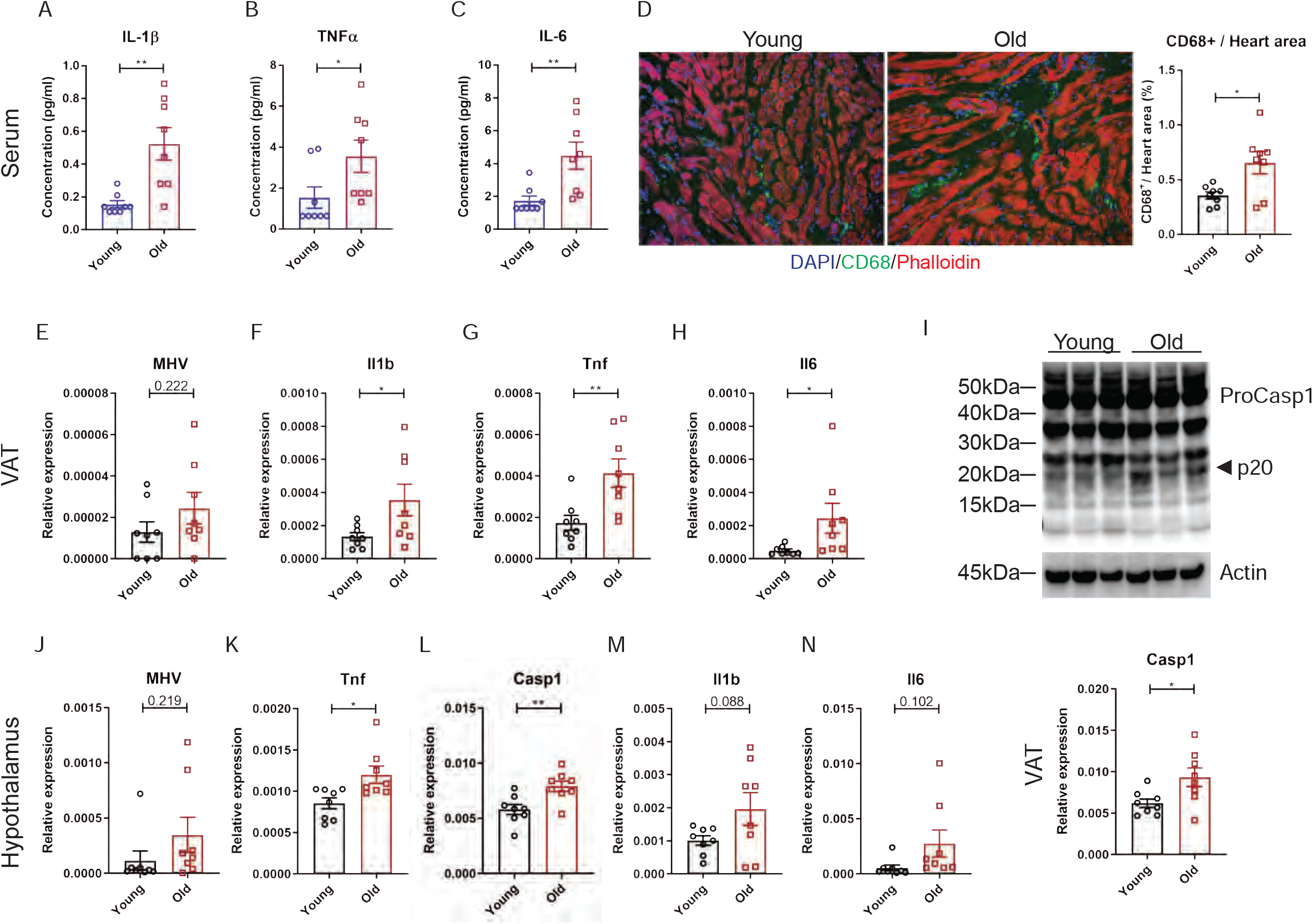
Aging induces systemic hyper-inflammatory response in A59 (mCoV) infection. **(A-C)** Serum level of inflammatory cytokines IL-1β (A), TNFα (B), and IL-6 (C) of young (6mo) and old (24mo) infected mice on day 8 post (PFU 7e2) infection. (**D)** Representative immunofluorescence analysis of CD68 expression, phalloidin, and DAPI in hearts isolated from young and old mice. CD68^+^ cells/heart area analysis was shown in right panel. (**E)** Quantification of MHV-A59 (mCoV) in visceral adipose tissue (VAT) of young and old infected mice by qPCR of A59 M protein. (**F-H)** qPCR analysis of *Il1b* (F), *Tnf* (G), and *Il6* (H) in VAT. (**I)** Immunoblot analysis of Caspase-1 cleavage showing higher inflammasome activation in VAT in aged mice post infection. Bottom panel indicates gene expression of *Casp1* in VAT. (**J)** Quantification of MHV-A59 (mCoV) in hypothalamus of young and old infected mice by qPCR. (**K-M)** Gene expression analysis of *Tnf* (K), *Casp1* (L), *Il1b* (M), and *Il6* (N) in hypothalamus of young and old mice 8 days post infection. Error bars represent the mean ± S.E.M. Two-tailed unpaired t-tests were performed for statistical analysis. * P < 0.05; ** P < 0.01.

Given that increased visceral adiposity is a risk factor for COVID-19 severity and expression of ACE2 is upregulated in adipocytes of obese and diabetic patients infected with SARS-CoV-2 (Kruglikov and Scherer, 2020), we next studied whether mCoV-A59 infection affects adipose tissue. Given the prevalence of obesity is 10% among younger adults aged 20-39, 45% among adults aged 40-59 years and 43% among older adults aged 60 and over (Hales et al., 2020) we investigated adipose tissue inflammation as a potential mechanism that contributes to infection severity in the aged. Interestingly, consistent with the prior findings that adipose tissue can harbor several viruses (Damouche et al., 2015), the mCoV-A59 RNA was detectable in VAT (Figure 3E). Despite similar viral loads, VAT of aged infected mice had significantly higher levels of the pro-inflammatory cytokines IL-1β, TNFα and IL-6 (Figure 3F-H). Moreover, compared to young mice, old animals infected with mCoV-A59 had increased Caspase-1 cleavage (p20 active heterodimer), a marker of inflammasome activation (Figure 3I and S2C). In addition, similar to SARS-CoV-2 invasiveness in CNS, the mCoV-A59 was detectable in the hypothalamus (Figure 3J). Compared to adults, the hypothalamus of aged infected mice showed increased expression of TNFα and Caspase-1 (Figure 3K and L) with no significant differences in IL-1β, IL-6 (Figure 3M and N) and Nlrp3 (Figure S2D). Infection in both young and aged mice caused significant increases in markers of astrogliosis and microglia activation (Figure S2E and F). Interestingly, mCoV-A59 reduced the mRNA expression of orexigenic neuropeptide-Y (NPY) (Figure S2E and F), consistent with the fact that infected mice display anorexia (Figure 1C and E). However, mCoV-A59 infection completely abolished the expression of pre-opiomelanocortin (POMC) in the hypothalamus, a transcript expressed by POMC neurons, which is involved in the control of the autonomic nervous system and integrative physiology. Therefore, further investigation will be necessary to test the involvement of the hypothalamus in the pathogenesis of COVID-19 and organ failure due to alterations in autonomic nervous system.

### Ketogenic diet mediated protection against A59 (mCoV) infection in aging is coupled to inflammasome deactivation

Given the switch from glycolysis to fatty acid oxidation reprograms the myeloid cell from pro-inflammatory to tissue reparative phenotype during infections (Ayres, 2020; Buck et al., 2017; Galván-Peña and O’Neill, 2014), we next investigated whether mCoV-A59-driven hyperinflammatory response in aging can be targeted through immunometabolic approaches. Hepatic ketogenesis, a process downstream of lipolysis that converts long-chain fatty acids into short chain β-hydroxybutyrate (BHB) as a preferential fatty acid fuel during starvation or glucoprivic states, inhibits the Nlrp3 inflammasome activation (Youm et al., 2015) and protects against influenza infection induced mortality in mice (Goldberg et al., 2019). Moreover, given our recent findings that ketogenesis inhibits inflammation and expands tissue resident ϒδ T cells (Goldberg et al., 2019) while SARS-CoV-2 infection in patients is associated with depletion of ϒδ T cells (Lei et al., 2020; Rijkers et al., 2020), we next tested whether elevating BHB by feeding a ketogenic diet (KD) protects against mCoV-A59-driven inflammatory damage in aged mice. We infected bone marrow derived macrophages (BMDMs) with mCoV-A59 *in vitro* in TLR4 (Figure 4A and S3A) and TLR1/2 primed cells (Figure 4B and S3B). Infection with mCoV-A59 caused robust activation of inflammasome as measured by cleavage of active IL-1β (p17) in BMDM supernatants (Figure 4A and B) as well as in cell lysates (Figure S3A and B). Given our prior findings that ketone metabolites specifically inhibits the NLRP3 inflammasome in response to sterile DAMPs such as ATP, ceramides, silica and urate crystals (Youm et al., 2015), we next tested whether BHB impacts inflammasome activation caused by mCoV-A59.

**Figure 4.**
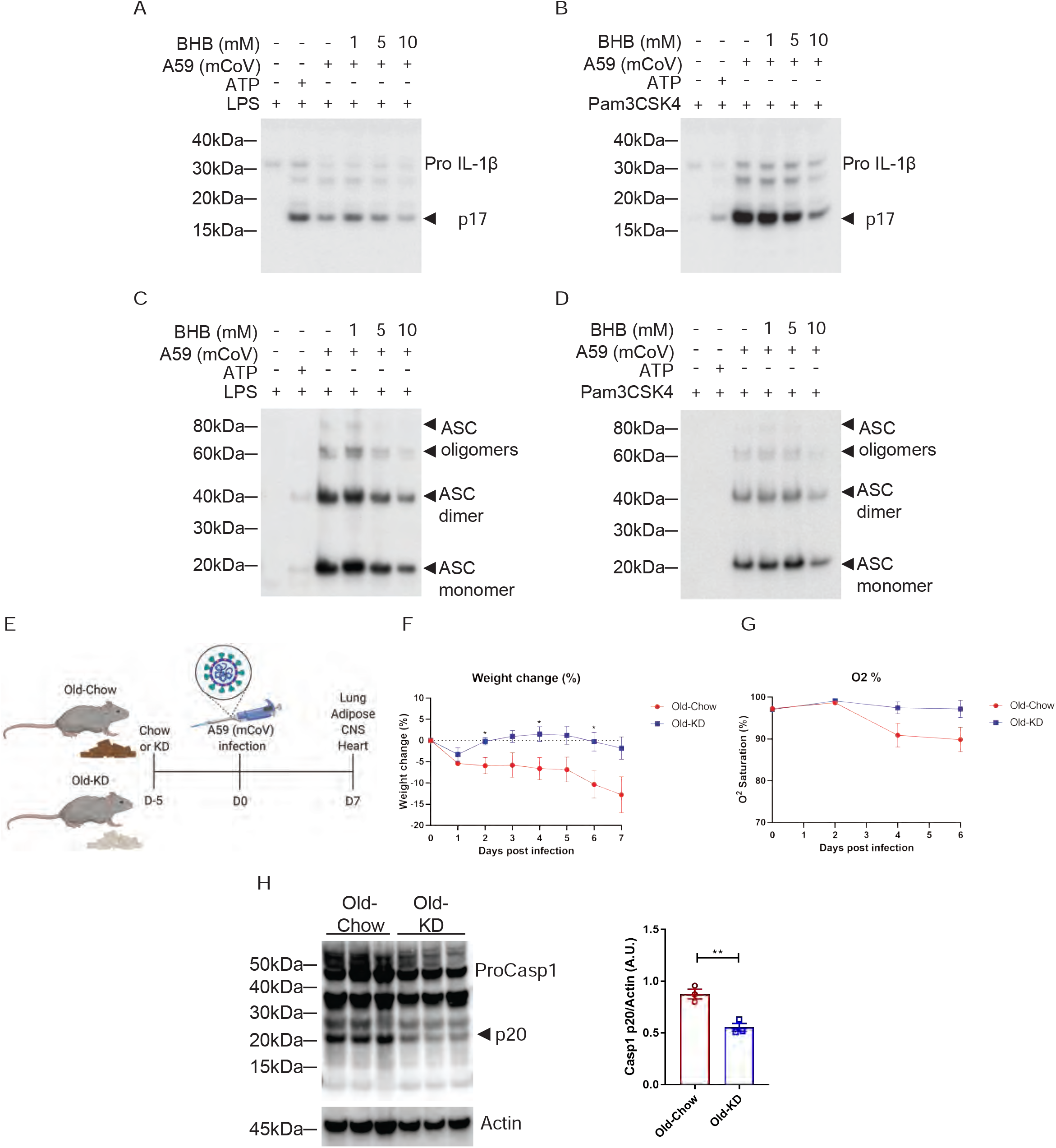
Ketogenic diet reduces inflammasome activation in A59 (mCoV) infected old mice. **(A-B)** Western blot analysis about pro and active cleaved p17 form of IL-1β from supernatant of A59 (mCoV) infected BMDMs co-treated with priming reagents such as LPS (A) and Pam3CSK4 (B), and BHB with indicated concentration. **(C-D)** Western blot analysis of ASC monomer, dimer, and oligomers from insoluble pellet of A59 (mCoV) infected BMDMs co-treated with priming reagents such as LPS (C) and Pam3CSK4 (D), and BHB with indicated concentration. **(E)** Schematic of A59 (mCoV) infection experiment with old mice (20-21 month) fed chow (Old-Chow, n=6) or ketogenic diet (Old-KD, n=5). The mice were provided with diet from 5 days before infection. After infection, the phenotype was evaluated until 7 days post infection. Weight change (%) (**F)**, and % O2 saturation (**G**) in old mice fed chow or KD. (**G)** Western blot analysis of Caspase-1 inflammasome activation in VAT of infected old-Chow and old-KD mice. Error bars represent the mean ± S.E.M. Two-tailed unpaired t-tests were performed for statistical analysis. * P < 0.05; ** P < 0.01.

Interestingly, BHB treatment reduced pro and active cleaved IL-1β (p17) in both conditions when protein level was measured in the supernatant (Figure 4A and B) and cell lysate (Figure S3A and B). Mechanistically, post mCoV-A59 infection, the BHB reduced the oligomerization of ASC, which is an adaptor protein required for the assembly of the inflammasome complex (Figure 4C and D). This data provides evidence that the ketone metabolite BHB can lower inflammation in response to coronavirus infection and deactivate the inflammasome. However, inflammasome activation is also required for mounting adequate immune response against pathogens including certain viruses. Therefore, we next investigated if induction of ketogenesis and ketolysis *in vivo* by feeding a diet rich in fat and low in carbohydrates that elevates BHB level impacts inflammasome and host defense against mCoV-A59 infection in aged mice.

Ketogenesis is dependent on hydrolysis of triglycerides and conversion of long chain fatty acids in liver into short chain fatty acid BHB that serves as primary source of ATP for heart and brain when glucose is limiting. Aging is associated with impaired lipid metabolism which includes reduced lipolysis that generates free fatty acids that are essential substrates for BHB production. Thus, it is unclear whether in context of severe infection and aging, if sufficient ketogenesis can be induced. To test this, the aged male mice (20-21 months old) were fed a KD or control diet for 5 days and then intranasally infected with mCoV-A59 (Figure 4E). Despite mCoV-A59’s known effects in causing hepatic inflammation (Navas et al., 2001), we observed that compared to chow fed animals, old KD fed mice achieved mild physiological ketosis between 0.6 to 1mM over the course of infection for one week (Figure S3C). Compared to chow fed animals, the mCoV-A59 infected mice fed KD displayed similar levels of food intake, blood glucose, core-body temperature, heart rate, and respiration (Figure S3D-I). Interestingly, mCoV-A59 infected KD fed mice were protected from infection-induced weight-loss and hypoxemia (Figure 4F and G). Importantly, consistent with *in vitro* data, KD feeding caused significant reduction in inflammasome activation in the VAT (Figure 4H), a major source of inflammation in aging. Together, these data show that KD lowers exuberant inflammasome activation in old mice and can potentially be therapeutically employed.

### Induction of ketogenic substrate switch inhibits systemic inflammation in aged mCoV-A59 infection

The elderly COVID-19 patients exhibit multi-organ failure with systemic viremia and inflammation. Therefore, we next investigated the impact of KD on the inflammatory response in lungs, adipose tissue and hypothalamus in old mice post mCov-A59 infection. Consistent with the improved clinical outcome and protection afforded by ketone bodies in infected mice, we found that KD-fed mice inoculated with mCoV-A59 had significantly reduced expression of pro-inflammatory cytokines IL-1β, TNFα and IL-6 in lung, VAT and hypothalamus (Figure 5A-C). Aging and mCoV-A59 increases inflammasome activation which is increasingly implicated in the pathogenesis of COVID-19 (Vijay et al., 2017; Youm et al., 2013). The SARS-CoV open reading frame 3a (ORF3a) and ORF8b activates the NLRP3 inflammasome (Siu et al., 2019) by inducing ER stress and lysosomal damage (Shi et al., 2019). Moreover, ability of bats to harbor multiple viruses including coronaviruses is due to splice-variants in the LRR domain of NLRP3 which prevents inflammasome mediated inflammatory damage (Ahn et al., 2019). Interestingly, in the aging mouse model of mCoV-A59 infection, KD significantly lowered NLRP3 and Caspase-1 mRNA in lung, VAT and hypothalamus (Figure 5D and E) and decreased myeloid cell infiltration in heart (Figure 5F). The ketogenesis in infected old mice did not affect the frequency of CD4, CD8 effector memory or macrophage subsets in lungs suggesting that reduction in pro-inflammatory cytokines was not a reflection of reduced infiltration of these cell types (Figure S4). Interestingly, we found that KD feeding rescued mCoV-A59-induced depletion of ϒδ T cell in lungs of aged mice (Figure 5G and S4A).

**Figure 5.**
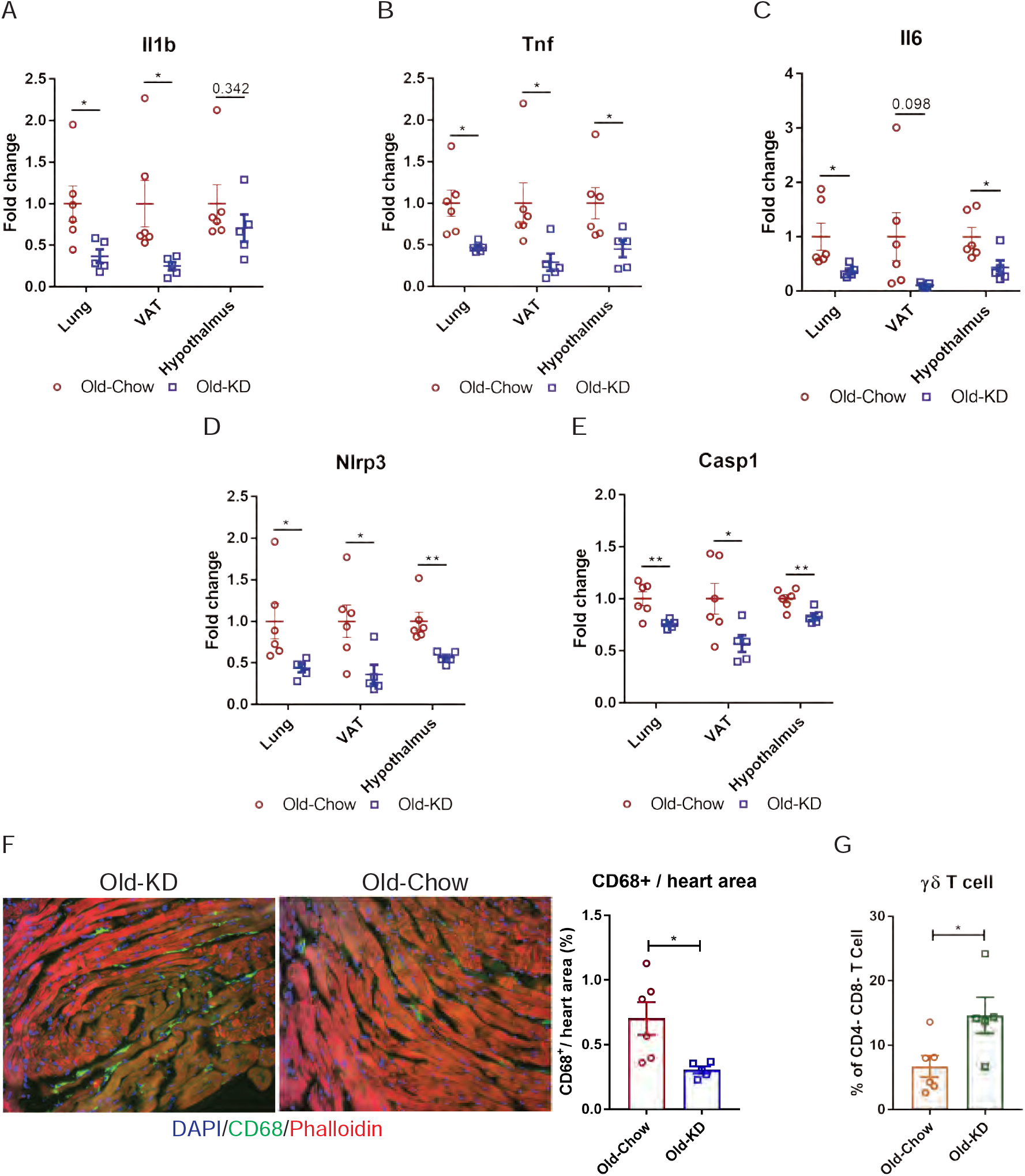
Ketogenic diet protects old mice from A59 (mCoV) infection by alleviation of inflammation. (**A-E)** Gene expression analysis of inflammatory cytokines including (A, B, C) and components of inflammasome (D, E) in lung, VAT, and hypothalamus of A59 (mCoV) infected old mice fed chow and old-KD mice. (**F)** Representative immunofluorescence analysis of CD68 expression, phalloidin, and DAPI in hearts isolated from old-Chow and old-KD mice (left) and CD68+ cells/heart area analysis in heart of infected old-Chow and old-KD mice (right). (**G)** Flow cytometry analysis of γδ T cell in lung of infected old-Chow and old-KD mice. Error bars represent the mean ± S.E.M. Two-tailed unpaired t-tests were performed for statistical analysis. * P < 0.05; ** P < 0.01.

### Ketogenesis induces protective γδ T cells and decreases myeloid cell subset in mCoV-A59 infected old mice

To determine the mechanism of ketogenesis-induced protection from mCoV-A59 driven inflammatory damage in aging, we next investigated the transcriptional changes in lung at the single-cell level. The scRNA sequencing of whole lung tissues (Figure 6A) found that KD feeding in old infected mice caused significant increase in goblet cells (Figure 6B), expansion of ϒδ T cells (Figure 6B) and significant decrease in proliferative cell subsets and monocyte populations (Figure 6B). Comparison with scRNA-seq of the lungs from young and old non-infected animals highlighted that only loss of proliferative myeloid cells was associated with the baseline aging process (Figure S5), while other age-specific changes in cellular subpopulations emerged as a result of interaction between virus and host. Moreover, the old mice showed reduced interferon responses, suggesting increased vulnerability to the viral infection (Figure S5). Interestingly, some of the most striking changes occurred in T cells, where ketogenesis led to a substantial increase in ϒδ but not αβ T cells (Figure 6C and Figure S6B). To understand whether expansion of ϒδ T cells was also accompanied by the changes in their regulatory programs, we sorted the lung ϒδ T cells from aged mice fed chow diet and KD and conducted bulk RNA sequencing to determine the mechanism of potential tissue protective effects of these cells in mCoV-A59 infection. We found that KD in aging significantly increased the genes associated with reduced inflammation (Figure 6D), increased lipoprotein remodeling and downregulation of TLR signaling, Plk1 and aurora B signaling pathways in ϒδ T cells (Figure 6E). Furthermore, RNA sequencing revealed that lung ϒδ T cells from ketogenic mCoV-A59 infected old mice displayed elevated respiratory electron transport and complex I biogenesis (Figure 6E). In addition, Golgi to ER retrograde transport and cell cycle are downregulated, suggesting the reduced activation status of ϒδ T cell (Figure 6E). This may indicate that ϒδ T cells expanded with KD are functionally more homeostatic and immune protective against mCoV-A59 infection.

**Figure 6.**
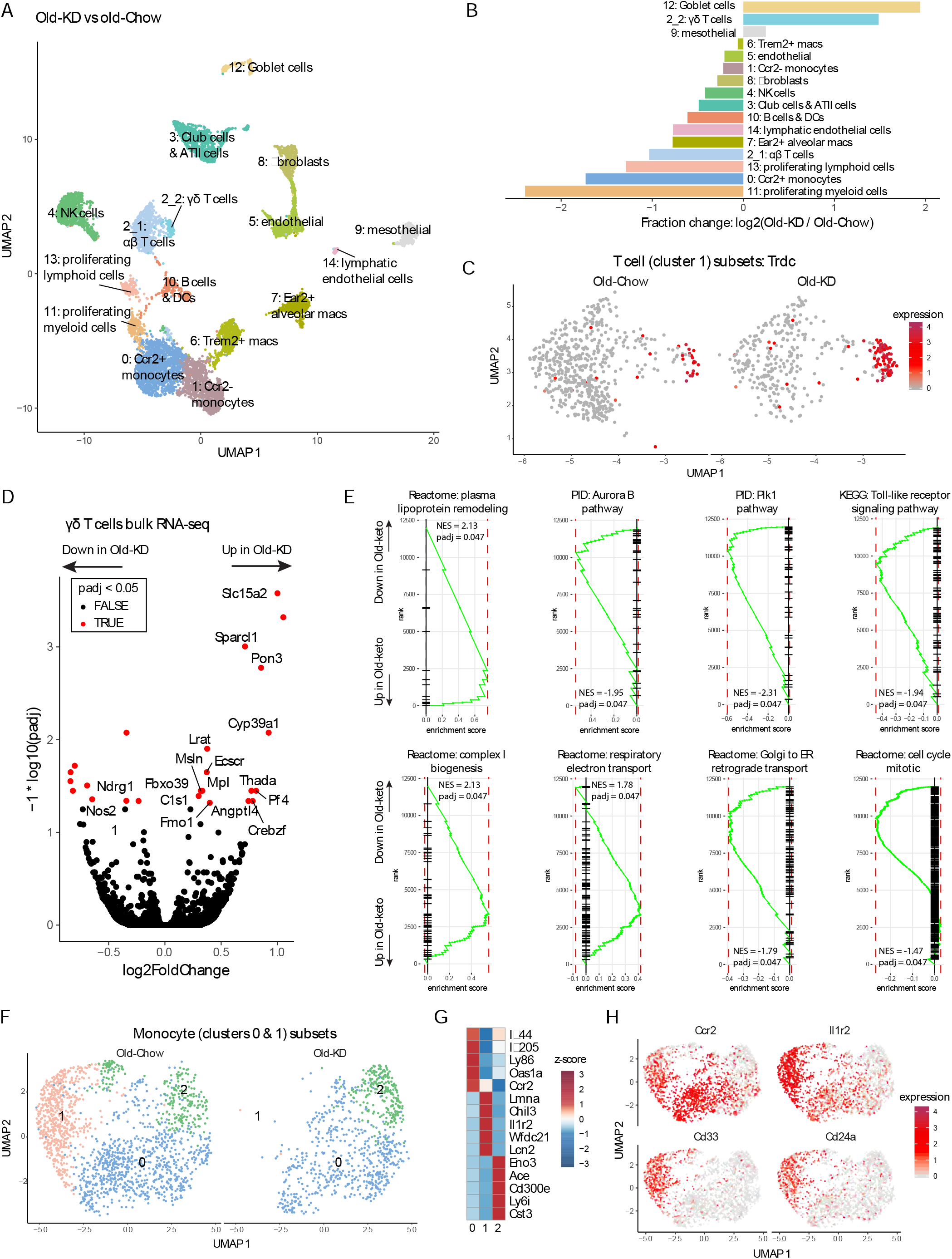
Ketogenesis induces protective γδ T cell expansion and inhibits myeloid cell activation in aged mice with mCoV infection. **(A)** UMAP plot of lung cells from Old-KD and Old-Chow samples as described in 3A. **(B)** Bar chart shows population fold-changes in relative abundance of each cluster. **(C)** Zoom into UMAP plot from 4A showing T cells (cluster 1) split by sample. Color represents expression of *Trdc*. **(D)** Volcano plot identifying significantly regulated genes (5% FDR) within sorted γσT cells from lungs of Old-KD and Old-Chow mice. Increase in expression corresponds to ketogenic diet-induced genes. **(E)** GSEA enrichment curve based on differential expression analysis results shown in 4D. **(F)** Monocyte cluster 0 and 1 from 4A were subset and analyzed separately. UMAP plot of monocytes split by samples. **(G)** Heatmap of normalized within row gene expression values of selected markers of three monocyte subsets. **(H)** UMAP as in 4F. Color represents expression of selected genes. For A-C, F-H, expression values were obtained by pooling data from Old-Chow and Old-KD samples (each containing n = 6 chow and n = 5 KD pooled biological samples into one technical sample for each diet).

Zooming in into monocyte subpopulation we observed three distinct monocyte clusters (Figure 6F), characterized by *Ifi44*, *Lmna*, and *Cd300e* respectively (Figure 6G). Strikingly, ketogenesis-induced change in the monocyte compartment was driven by a loss of cluster 1 (characterized by high levels of *Chil3*, *Lmna*, *Il1r2*, *Lcn2*, *Cd33*, *Cd24a*, Figure 6H and Figure S6D and E). In addition, the loss of monocyte subpopulation was observed in cells with low interferon-response further suggesting the immune protective response induction post ketogenesis in infected mice. This is an intriguing finding that is consistent with recent observations that dietary interventions can impact plasticity of the monocyte pool in both mouse and human (Collins et al., 2019; Jordan et al., 2019).

## DISCUSSION

Immune-senescence exemplified by inflammasome-mediated basal activation of myeloid cells, expansion of pro-inflammatory aged B cells, impaired germinal center and antibody responses together with thymic demise and restriction of T cell repertoire diversity all contribute to increased risk of infections and vaccination failures in elderly (Akbar and Gilroy, 2020; Frasca et al., 2020; Goldberg and Dixit, 2015; Goronzy and Weyand, 2019). It is likely that multiple mechanisms partake in aging-induced mortality and morbidity to SARS-CoV-2. However, study of immunometabolic mechanisms that control aberrant inflammatory response in elderly COVID-19 patients are hindered due to lack of availability of an aging mouse model of disease that recapitulates the key features of SARS-CoV-2 immunopathology and multi-organ inflammation. Despite the severity of this viral infection, it is currently unclear what underlies the symptom diversity and the mortality of this pandemic. Epidemiological data strongly support that elderly and aged individuals with late-onset chronic diseases—including diabetes, obesity, heart conditions, pulmonary dysfunctions and cancer—present a much higher disease severity compared to young healthy adults (Cai et al., 2020; Chen et al., 2020). These observations suggest that it is the vulnerability of the various tissues that occur in these chronic conditions that predispose elderly to develop severe forms of COVID-19.

Rodent CoVs are natural, highly contagious pathogens of mice and rats (Compton et al., 2003; Compton et al., 1993). They are efficient and safe platforms for recapitulating COVID-19 and examining factors and interventions that impact disease. These models enable basic and translational COVID-19 studies by minimizing studies requiring SARS-CoV-2 infection, thereby conserving and reserving limited BSL-3 space for studies using the most promising candidates in a homologous SARS-CoV-2 model. Among the diverse rodent CoVs, Mouse Hepatitis Virus (MHV) is a collection of mouse CoV strains which have clinical diseases ranging from clinically silent (enterotropic) to mortality (polytropic/respiratory tropic). Of particular interest for COVID-19 are the strains of MHV that are respiratory tropic (Yang et al., 2014). Given, these advantages, MHV mCoV-A59 infection in C57BL/6 mice can be a powerful tool to rapidly study the disease as well as test therapeutic interventions. We demonstrate that compared to all reported models (Dinnon et al., 2020; Hassan et al., 2020; Israelow et al., 2020; Jiang et al., 2020; Sun et al., 2020; Winkler et al., 2020), MHV mCoV-A59 infection recapitulates severe features of COVID-19 that includes, up to 30% weight-loss, sickness behavior exemplified by anorexia, loss of oxygen saturation, lung pathology including neutrophilia, monocytosis, loss of γδ T cells, lymphopenia, increase in circulating pro-inflammatory cytokines, hypothalamic, adipose and cardiac inflammation and inflammasome activation. Importantly, LD0 dose of MHV mCoV-A59 induces 100% lethality in 2 year old male mice, suggesting that this model allows investigation of COVID-19 relevant immunometabolic mechanisms that control disease development and severity with aging.

Mechanistically, NLRP3 inflammasome has been demonstrated to be an important driver of aging-induced chronic inflammation and organ damage (Bauernfeind et al., 2016; Camell et al., 2017; Youm et al., 2013). COVID-19 patients have inflammasome dependent pyroptosis and increase in IL-18 (Lucas et al., 2020; Zhou et al., 2020). Consistent with the hypothesis that aging may exacerbate inflammasome activation in SARS-CoV-2 infection, our data demonstrates that *in vivo*, mCoV infection increases NLRP3 inflammasome mediated inflammation. Emerging evidence demonstrates that ability of bats to harbor multiple viruses including coronaviruses is due to splice-variants in the LRR domain of NLRP3 which prevents inflammasome mediated inflammatory damage (Ahn et al., 2019). Furthermore, recent study shows that MHV-A59 also activates the NLRP3 inflammasome *in vitro* bone marrow derived macrophages (Zheng et al., 2020). In addition severe cases of COVID-19 are accompanied with dyregulation of monocyte populations with increased level of S100A8/A9 or calprotectin (Schulte-Schrepping et al., 2020; Silvin et al., 2020), which can prime and induce the inflammasome activation (Goldberg et al., 2017). These data underscore that enhanced innate immune tolerance mediated by inflammasome deactivation maybe an important strategy against COVID-19.

The integrated immunometabolic response (IIMR) is critical in regulating the setpoint of protective versus pathogenic inflammatory response (Lee and Dixit, 2020). The IIMR involves sensing of nutrient balance by neuronal (sympathetic and sensory innervation) and humoral signals (e.g. hormones and cytokines) between the CNS and peripheral tissues that allow the host to prioritize storage and/or utilize substrates for tissue growth, maintenance and protective inflammatory responses. Peripheral immune cells, both in circulation and those residing within tissues, are subject to regulation by the metabolic status of the host. Ketone bodies, BHB and acetoacetate are produced during starvation to support the survival of host by serving as an alternative energy substrate when glycogen reserves are depleted (Newman and Verdin, 2017). Classically, ketone bodies are considered essential metabolic fuels for key tissues such as the brain and heart (Puchalska and Crawford, 2017; Veech et al., 2017). However, there is increasing evidence that immune cells can also be profoundly regulated by ketone bodies (Youm et al, 2015, Goldberg et al 2020). For example, stable isotope tracing revealed that macrophage oxidation of liver-derived AcAc was essential for protection against liver fibrosis (Puchalska et al., 2019). Given our past findings that ketone bodies inhibit NLRP3 inflammasome activation induced by sterile DAMPs, we next hypothesized that coronavirus mediated inflammasome activation and disease severity in aging could be improved by BHB driven improved metabolic efficiency and NLRP3 deactivation. In support of this hypothesis, we found that BHB inhibits the mCoV-A59 induced NLRP3 inflammasome assembly and KD reduces caspase-1 cleavage as well as decreases gene expression of inflammasome components. We next investigated the mechanism of protection elicited by KD that is relevant to aging. Interestingly, scRNA sequencing analyses of lung homogenates of old mice fed KD revealed robust expansion of immunoprotective γδ T cells, which are reported to decline in COVID-19 patients (Lei et al., 2020; Rijkers et al., 2020). The KD activated the mitochondrial function as evidenced by enhanced complex-1 biogenesis and upregulation of ETC in immunoprotective γδ T cells. Moreover, the KD feeding blocked infiltration of pathogenic monocyte subset in lungs that has high S100A8/9 and low interferon expression.

Our data shows that the mCoV-A59 murine model offers an efficient and biosafety level-2 platform for recapitulating COVID-19 to test mechanism of age-related immune decline and can thereby fast-track testing of interventions that impact disease outcome. Our findings assumes strong clinical significance as recent studies, demonstrate that γδ T cells were severely depleted in COVID-19 patients in two highly variable cohorts and disease progression was correlated with near ablation of Vγ9Vδ2 cells that are dominant subtype of circulating γδ T cells (Laing et al., 2020). Taken together these data demonstrate that a ketogenic immunometabolic switch protects against mCoV-A59 driven COVID in mice and this anti-inflammatory response in lung is coupled with reduction of inflammasome activation, restoration of protective ϒδ T cells and remodeling of the pool of the inflammatory monocytes. Finally, our results suggest that acutely switching infected or at-risk elderly patients to a KD may ameliorate COVID-19 and, therefore, is a relatively accessible and affordable intervention that can be promptly applied in most clinical settings.

## LIMITATIONS

Mouse remains imperfect to model human biology and disease. Instead of following the current approaches to make human SARS-CoV-2 amenable to infecting an unnatural murine host with unknown biological compatibility, we focused on natural mouse hepatitis virus (MHV)-A59 because like SARS-CoV-2, it belongs to the family of ARDS-related beta coronaviruses that are highly homologous. However, the obvious limitation of the model is that MHV-A59 is not SARS-CoV-2 virus. Although, both these beta coronaviruses display high degree of homology, MHV-A59 uses CEACAM1 instead of ACE2 for binding and infectivity. However, cellular expression of CEACAM1 is similar to ACE2 in humans and similarity of viral ORFs offer significant advantages in studying tissue responses. Like most models of disease, this study shows that not all features of SARS-CoV-2 infection seen in humans are seen in mice. This includes lack of development of fever instead mice become hypothermic. Also, despite monocyte infiltration in heart, the aged mice did not die due to cardiac failure and displayed normal heart rate. mCoV-A59 virus induced hypothalamic inflammation and led to anorexia that included reduction in orexigenic NPY but almost complete loss of POMC gene expression. It remains unclear, if this alters POMC derived peptides including melanocortins and endogenous opioids peptides, in addition these data suggest potential dysregulation of autonomic nervous system which can play a role in organ failure post infection. In terms of mechanism, our data shows that the inflammasome is activated in infection and KD-induced protection is associated with NLRP3 inflammasome deactivation. Future studies are required to test if aged NLRP3 deficient mice are protected from mCoV-A59 or if absence of γδ T cell increases mortality.

## ACKNOWLEDGMENTS

SR is supported by American Federation of Aging Research (AFAR) postdoctoral fellowship award. The Dixit lab is supported in part by NIH grants P01AG051459, AR070811 and Cure Alzheimers Fund. The Wang lab is supported in part by NIH grant 1K08AI128745 and by gifts from the Knights of Columbus, G. Harold and Leila Y. Mathers Charitable Foundation, and the Ludwig Family Foundation. We also thank the Yale Center on Genomic Analysis (YCGA) for RNA-seq studies and Genentech Inc for providing the anti-caspase-1 antibody.

## AUTHOR CONTRIBUTIONS

SR performed experiments, data analysis, and prepared the manuscript. IS and MA performed analyses of data from single cell RNA-sequencing and bulk RNA-sequencing experiments. YY, and TD performed experiments regarding mouse tissue processing, and qPCR. HQ and BH each performed the infections, phenotyping and plaque assay. XZ, YS and SF analyzed the impact of infection on heart. TLH, YY designed and executed electron microscopy, CB performed the histological analyses. MOD designed experiments and analyzed hypothalamic inflammation. KMK designed lung immune cell analyses experiments. CF, YS, KMK, TLH, MOD, and MNA provided critical reviews and helped in organization of experiments and interpretation of data. AW and VDD conceived the project and helped with data interpretation and manuscript preparation.

## STAR★METHODS

### KEY RESOURCES TABLE

### RESOURCE AVAILABILITY

#### Lead Contact

Further information and requests for resources and reagents should be directed to and will be fulfilled by the Lead Contact, Vishwa Deep Dixit.

#### Materials Availability

This study did not generate new unique reagents.

#### Data and Code Availability

The single cell RNA-sequencing and bulk RNA-sequencing data has been uploaded to Gene Expression Omnibus (GSE155346 and GSE155347) respectively.

### EXPERIMENTAL MODEL AND SUBJECT DETAILS

#### Mice

All mice used in this study were C57BL/6 mice. Old mice (20-24 month old) were received from NIA, maintained in our laboratory. Young mice (2-6 month old) were from NIA or purchased from Jackson Laboratories or bred in our laboratory. The mice were housed in specific pathogen-free facilities with free access to sterile water through Yale Animal Resources Center. Mice were fed a standard vivarium chow (Harlan 2018s) or a ketogenic diet (Envigo, TD.190049) for indicated time. The mice were housed under 12 h light/dark cycles. All experiments and animal use were conducted in compliance with the National Institute of Health Guide for the Care and Use of Laboratory Animals and were approved by the Institutional Animal Care and Use Committee (IACUC) at Yale University.

### METHOD DETAILS

#### Viral infection

MHV-A59 was purchased from Bei resources (NR-43000) and grown in BV2 cells. Mice were anesthetized by intraperitoneal injection of ketamine/xylazine. 700 or 7000 PFU of MHV-A59 was delivered in 40ul PBS via intranasal inoculation. Vital signs were measured before and after infection. Arterial oxygen saturation, breath rate, heart rate, and pulse distention were measured in conscious, unrestrained mice via pulse oximetry using the MouseOx Plus (Starr Life Sciences Corp.).

#### Electron Microscopy

Lungs were fixed in 10% formaldehyde, osmicated in 1% osmium tetroxide, and dehydrated in ethanol. During dehydration, 1% uranyl acetate was added to the 70% ethanol to enhance ultrastructural membrane contrast. After dehydration, the lungs were embedded in Durcupan and ultrathin sections were cut on a Leica Ultra-Microtome, collected on Formvar-coated single-slot grids, and analyzed with a Tecnai 12 Biotwin electron microscope (FEI).

#### Histology and Immunohistochemistry

H&E and MSB staining of lung tissues were performed on sections of formalin-fixed paraffin-embedded at the Comparative Pathology Research core at Yale School of Medicine. For immunohistochemistry, the hearts were harvested from MHV-A59 infected mice, fixed in 4% PFA overnight and embedded in OCT after dehydration with 30% sucrose and serial sections of aortic root were cut at 6 μm thickness using a cryostat. Sections were incubated at 4°C overnight with CD68 (Serotec; #MCA1957) and Alexa Fluor™ 594 Phalloidin (ThermoFisher, A12381) after blocking with blocker buffer (5% Donkey Serum, 0.5% BSA, 0.3% Triton X-100 in PBS) for 1 hour at RT, followed by incubation with Alexa Fluor secondary antibody (Invitrogen, Carlsbad, CA) for 1 hour at RT. The stained sections were captured using a Carl Zeiss scanning microscope Axiovert 200M imaging system and images were digitized under constant exposure time, gain, and offset. Results are expressed as the percent of the total plaque area stained measured with the Image J software (ImageJ version 1.51).

#### Plaque assay

L2 cell (1.5 mL of 6×10^5^ cells/mL) were seeded on 6 well plates (Corning, 3516) in supplemented DMEM and allowed to adhere overnight. Tissue samples were homogenized in unsupplemented DMEM, and spun down at 2000 rpm for 5 min. Supernatant was serially diluted and 200 μL of each sample was added to aspirated L2 cells in 6 well plates. Plates were agitated regularly for 1 hour before adding overlay media consisting of 1 part 1.2% Avicel and 1 part 2X DMEM (Thermo Fisher, 12800) supplemented with 4% FBS (Thermo Fisher, A3840301), Penicillin-Streptomycin (Gibco, 15140122), MEM Non-Essential Amino Acids Solution (Gibco, 11140050), and HEPES (Gibco, 15630080). After a four-day incubation, cells were fixed in 10% formaldehyde (Sigma Aldrich, 8187081000) diluted with PBS for 1 hour. Cells were then stained in 1% (w/v) crystal violet (Sigma Aldrich, C0775) for 1 hour, washed once in distilled water, and then quantified for plaque formations.

#### Multiplex Cytokine Analyses

Serum cytokine and chemokine level was measured by ProcartaPlex multiplex assay (Thermo Fisher Scientific). Assay was prepared following manufacture’s instruction. 25 μl of collected serum from each mice in this study was used. Customized assay including IL-1β, TNFα, IL-6, MCP-1, and MIP-1β was used. Luminex xPONENT system was used to perform the assay.

#### qPCR

To extract and purify RNA from tissues, RNeasy Plus micro kit (Qiagen) and Direct-zol™ RNA Miniprep Plus kit (Zymo Research) were used according to manufacturer’s instructions. cDNA was synthesized with isolated RNA using iScript cDNA synthesis kit (Bio-Rad). To quantify amount of mRNA, real time quantitative PCR (qPCR) was done with the synthesized cDNA, gene specific primers, and Power SYBR Green detection reagent (Thermo Fischer Scientific) using the LightCycler 480 II (Roche). Analysis was done by ΔΔCt method with measured values from specific genes, the values were normalized with *Gaphd* gene as an endogenous control.

#### BMDM culture and in vitro viral infection

Bone marrow derived macrophage was cultured by collecting mouse femurs and tibias in complete collecting media containing RPMI (Thermo Fischer Scientific), 10% FBS (Omega Scientific), and 1% antibiotics/antimycotic (Thermo Fischer Scientific). Using needle and syringe, bone marrow was flushed into new complete media, followed by red blood cells lysis by ACK lyses buffer (Quality Biological). In 6 well plate, the collected cells were seeded to be differentiated into macrophages incubated with 10 ng/ml M-CSF (R&D) and L929 (ATCC) conditioned media. Cells were harvested on day 7, and seeded as 1×10^6^ cell/well in 24 well plate for experiments. To infect BMDM, MHV-A59 was incubated with BMDM as a MOI 1 (1:1) for 24 hour. For inflammasome activation, LPS (1ug/ml) or Pam3CSK4 (1ug/ml) were pre-treated with or without BHB (1, 5, 10 mM) for 4 hour before MHV-A59 infection for 24 hour or ATP (5mM) treatment for 1 hour.

#### Western Blotting

To prepare samples for western blotting, tissues were snap frozen in liquid nitrogen. RIPA buffer with protease inhibitors were used to homogenize the tissues. After cell supernatant was collected, cells were harvested by directly adding RIPA buffer on cell culture plate. After quantification of protein amount by the DC protein assay (Bio-Rad), same amount of protein was run on SDS-PAGE gel followed by transferring to nitrocellulose membrane. Specific primary antibodies and appropriate secondary antibodies (Thermo Fisher Scientific) were used to probe blots and bands were detected by ECL Western Blotting Substrate (Pierce). The following primary antibodies were used for experiments. Antibodies to Caspase-1 (1:250, Genentech), β-actin (1:1,000, 4967L; Cell Signaling), IL-1β (1:1000, GTX74034, GeneTex), and ASC (1:1000, AG-25B-0006, AdipoGen) were used.

#### ASC oligomerization assay

To detect ASC oligomers, cells were harvested in NP-40 lysis buffer which contains 20mM HEPES-KOH (pH 7.5), 150 mM KCl, 1% NP-40, 0.1 mM PMSF, and protease inhibitors. The cells in lysis buffer were incubated on ice for 15 min, and centrifuged at 6,000 rpm at 4°C for 10 min. Supernatant was collected and kept for cell lysate western blotting. The pellet was vortexed with 1 ml of NP-40 lysis buffer, and centrifuged at 6,000 rpm at 4°C for 10 min. The pellet was incubated with 50ul of NP-40 lysis buffer and 1 ul of 200 mM DSS (disuccinimidyl suberate) for 30 min at room temperature, then centrifuged at 6,000 rpm at 4°C for 10 min. The pellet with SDS sample buffer and reducing reagent was loaded for western blotting.

#### Flow Cytometry

Lung was digested in RPMI (Thermo Fisher) with 0.5mg/ml Collagenase I (Worthington) and 0.2mg/ml DNase I (Roche) for 1 hour. Digested lung tissues were minced through 100 μm strainer. Spleen was directly minced through 100 μm strainer. Minced tissues were additionally filtered with 40 μm strainer after red blood cell lysis by ACK lysing buffer (Quality Biological). After incubation with Fc Block CD16/32 antibodies (Thermo Fisher Scientific), the cells from lung and spleen were further incubated with surface antibodies for 30 min on ice in the dark. Washed cells were stained with LIVE/DEAD™ Fixable Aqua Dead Cell Stain Kit (Thermo Fisher Scientific). BD LSRII was used for Flow Cytometry and results were analyzed by FlowJo software. The following antibodies were used for flow cytometry analysis to detect CD4 T cell, CD8 T cell, γδ T cell, neutrophil, eosinophil, and macrophage: CD45-BV711, MerTK-FITC, CD64-BV605, F4/80-eFluor450, Ly6C-PerCP-Cy5.5, CD11c-APC, CD169-PE, CD86-PE-Cy7, CD3-BV605, CD4-PE-Cy7, CD8-eFluor450, TCR γ/δ-PE, Ly6G-APC, SiglecF-PerCP-Cy5.5, CD62L-PerCP-Cy5.5, CD44-APC-Cy7.

#### Single-cell RNA sequencing

Lung cells were prepared as mentioned above for flow cytometry and equal amount of cells were pooled as indicted in the experiments. Single-cell RNA sequencing libraries were prepared at Yale Center for Genome Analysis following manufacturer’s instruction (10x Genomics). NovaSeq6000 was used for sequencing library read.

##### Alignment, barcode assignment and unique molecular identifier (UMI) counting

The Cell Ranger Single-Cell Software Suite (v3.0.2) (available at https://support.10xgenomics.com/single-cell-gene-expression/software/pipelines/latest/what-is-cell-ranger) was used to perform sample demultiplexing, barcode processing, and single-cell 3’ counting. Cellranger mkfastq was used to demultiplex raw base call files from the NovaSeq6000 sequencer into sample-specific fastq files. Subsequently, fastq files for each sample were processed with cellranger counts to align reads to the mouse reference (version mm10-3.0.0) with default parameters.

##### Preprocessing analysis with Seurat package

For the analysis, the R (v3.5.0) package Seurat (v3.1.1) (Butler et al., 2018) was used. Cell Ranger filtered genes by barcode expression matrices were used as analysis inputs. Samples were pooled together using the merge function. The fraction of mitochondrial genes was calculated for every cell, and cells with high (>5%) mitochondrial fraction were filtered out. Expression measurements for each cell were normalized by total expression and then scaled to 10,000, after that log normalization was performed (NormalizeData function). Two sources of unwanted variation: UMI counts and fraction of mitochondrial reads – were removed with ScaleData function. For both datasets platelet clusters as well as a cluster of degraded cells (no specific signature and low UMI count) were removed and data was re-normalized without them. In case of Old-Keto and Old-Chow dataset we additionally removed neutrophils, doublets, and red blood cells.

##### Dimensionality reduction and clustering

The most variable genes were detected using the FindVariableGenes function. PCA was run only using these genes. Cells are represented with UMAP (Uniform Manifold Approximation and Projection) plots. We applied RunUMAP function to normalized data, using first 15 PCA components. For clustering, we used functions FindNeighbors and FindClusters that implement SNN (shared nearest neighbor) modularity optimization-based clustering algorithm on top 20 PCA components using resolution of 0.3 for both datasets.

##### Identification of cluster-specific genes and marker-based classification

To identify marker genes, FindAllMarkers function was used with likelihood-ratio test for single cell gene expression. For each cluster, only genes that were expressed in more than 10% of cells with at least 0.1-fold difference (log-scale) were considered. For heatmap representation, mean expression of markers inside each cluster calculated by AverageExpression function was used.

##### Single cell RNA-seq differential expression

To obtain differential expression between clusters, MAST test was performed via FindMarkers function on genes expressed in at least 1% of cells in both sets of cells, and p-value adjustment was done using a Bonferroni correction (Finak et al., 2015). Pathway analysis was performed using clusterProfiler package (v3.12.0) (Yu et al., 2012) with Hallmark gene sets from MSigDB. Significantly different genes were used (padj < 0.05) if percent difference between conditions (|pct.1 – pct.2|) was over 1%. To visualize pathway expression for each cell z-scores of all pathway genes were averaged.

##### Cell subsets analysis

To separate αβT cells and γσT cells we subset raw values of T cell (cluster 2), normalized and clustered corresponding data separately as described above with clustering resolution 0.2. Obtained subclusters were projected on the original UMAP, splitting cluster 2 in 2_1 (αβT cells) and 2_2 (γσT cells). Monocyte clusters 0 and 1 were subset and re-analyzed in the same manner with clustering resolution 0.3. UMAP was recalculated for monocytes only as shown in **Figure 6F**.

#### Bulk RNA sequencing of sorted γδ T cells

Cells from lung were prepared as mentioned above for flow cytometry and γδ T cell was sorted by flow cytometry (live CD45+ CD3+ CD4-CD8-TCR γ/δ+). RNA was isolated from sorted cells using RNeasy Plus micro kit (Qiagen). Quality checked RNA was used for RNA sequencing library preparation at Yale Center for Genome Analysis following manufacturer’s instruction (Illumina). NovaSeq6000 was used for sequencing library read. Fastq files for each sample were aligned to the mm10 genome (Gencode, release M25) using STAR (v2.7.3a) with the following parameters: STAR --genomeDir $GENOME_DIR -- readFilesIn $WORK_DIR/$FILE_1 $WORK_DIR/$FILE_2 --runThreadN 12 -- readFilesCommand zcat --outFilterMultimapNmax 15 --outFilterMismatchNmax 6 -- outReadsUnmapped Fastx --outSAMstrandField intronMotif --outSAMtype BAM SortedByCoordinate --outFileNamePrefix ./$ (Dobin et al., 2013). Quality control was performed by FastQC (v0.11.8), MultiQC (v1.9) (Ewels et al., 2016), and Picard tools (v2.21.6). Quantification was done using htseq-count function from HTSeq framework (v0.11.2): htseq-count -f bam -r pos -s no -t exon $BAM $ANNOTATION > $OUTPUT (Anders et al., 2014). Differential expression analysis was done using DESeq function from DeSeq2 package (Love et al., 2014) (v1.24.0) with default settings. Significance threshold was set to adjusted p-value < 0.05. Gene set enrichment analysis via fgsea R package (Sergushichev, 2016) (v1.10.0) was used to identify enriched pathways and plot enrichment curves.

### QUANTIFICATION AND STATISTICAL ANALYSIS

#### Statistical Analysis

To calculate statistical significance, two-tailed Student’s t test was used. Level of significance was indicated as follow. *p < 0.05; **p < 0.005; ***p < 0.001; ****p < 0.0001 respectively. All statistical tests used 95% confidence interval and normal distribution of data was assumed. Biological replication numbers for each experiment were indicated in each figure and figure legend. Data were shown as mean ± S.E.M. GraphPad Prism software was used for all statistical tests to analyze experimental results.

## SUPPLEMENTAL INFORMATION TITLES AND LEGENDS

**Figure S1. Characterization of immune cell population in young and old mice infected with A59 (mCoV), Related to Figure 1. (A)** Plaque assay of lung from infected young and old mice. **(B)** Flow cytometry analysis result of CD4/CD8 T cell ratio in lung of infected young and old mice. **(C)** Representative flow cytometry gating plot to analyze T lymphocytes in lung of infected young and old mice. Single cell gating was not shown in the plot. (**D)** Flow cytometry analysis result of γδ T cell in spleen of infected young and old mice. **(E)** Representative flow cytometry gating plot to analyze myeloid cells in lung of infected young and old mice. Single cell gating was not shown in the plot. (**F)** Flow cytometry analysis result of eosinophil in lung of infected young and old mice. **(G-H)** Flow cytometry analysis results of alveolar macrophage (AM) (G), and interstitial macrophage (IM) (H) population in lung of infected young and old mice. Error bars represent the mean ± S.E.M. Two-tailed unpaired t-tests were performed for statistical analysis. *** P < 0.001.

**Figure S2. Inflammatory response in young and old mice infected with A59 (mCoV), Related to Figure 3. (A, B)** Serum level of inflammatory chemokines of young and old infected mice. Analysis results of MCP-1 (A), and MIP-1β (B) were shown. **(C)** Western blot analysis of Caspase-1 inflammasome activation in VAT of uninfected young and old mice **(D)** Gene expression analysis of Nlrp3 in hypothalamus of young and old infected mice. **(E, F)** Gene expression analysis in hypothalamus of young uninfected and young infected mice (E), and old uninfected and old infected mice (F). Error bars represent the mean ± S.E.M. Two-tailed unpaired t-tests were performed for statistical analysis. * P < 0.05; ** P < 0.01; *** P < 0.001; **** P < 0.0001.

**Figure S3. Protective effect of BHB in BMDM against A59 (mCoV) and phenotype of A59 (mCoV) infected old mice fed chow or KD, Related to Figure 4. (A-B)** Western blot analysis about pro and active cleaved p17 form of IL-1β from cell lysate of A59 (mCoV) infected BMDMs co-treated with priming reagents such as LPS (A) and Pam3CSK4 (B), and BHB with indicated concentration. **(C-F)** Physiological phenotype of old-Chow and old-KD infected mice. Analysis results of β-hydroxybutyrate (BHB) level (C), food intake per day (D), glucose level (E), and core body temperature (F) were shown until 7 days post infection. **(G-I)** Measurement of vital signs of old-Chow and old-KD infected mice. Analysis results of heart rate (G), breath rate (H), and pulse distention (I) were indicated. Error bars represent the mean ± S.E.M. Two-tailed unpaired t-tests were performed for statistical analysis. * P < 0.05; ** P < 0.01; **** P < 0.0001.

**Figure S4. Immune cell population profile in lung of infected old mice provided with chow or ketogenic diet, Related to Figure 5. (A)** Representative flow cytometry gating plot to analyze T lymphocytes in lung from infected old-Chow and old-KD mice. Single cell gating was not shown in the plot. **(B-F)** Flow cytometry analysis of T lymphocytes and neutrophil in lung of infected old-Chow and old-KD mice. Analysis results of CD4 T cell (B), CD8 T cell (C), CD4/CD8 T cell ratio (D), γδ T cell (E), and neutrophil (F). **(G)** Representative flow cytometry gating plot to analyze myeloid cells in lung from infected old-Chow and old-KD mice. Single cell gating was not shown in the plot. **(H-L)** Flow cytometry analysis of myeloid cells in lung of infected old-Chow and old-KD mice. Analysis results of CD64^+^ MerTK^+^ cell (H), Ly6C^hi^ cell (I), F4/80+ Ly6C^lo^ cell (J), alveolar macrophage (K), and interstitial macrophage (L). **(M-T)** Sub-population analysis of T cells and macrophages in lung of infected old-Chow and old-KD mice by flow cytometry analysis. CD44+ CD62L-(M) and CD44-CD62+ (N) cell population in CD4 T cells. CD44+ CD62L-(O) and CD44-CD62+ (P) cell population in CD8 T cells. CD44+ CD62L-(Q) and CD44-CD62+ (R) cell population in γδ T cells. CD86+ alveolar macrophage (S) and interstitial macrophage (T). Error bars represent the mean ± S.E.M. Two-tailed unpaired t-tests were performed for statistical analysis. * P < 0.05.

**Figure S5. Single cell RNA-sequencing analysis of lung from young and old A59 (mCoV) infected mice, Related to Figure 6. (A)** UMAP plot of lung cells from young and old samples as described in **1A**. **(B)** Heatmap of normalized gene expression values of selected genes to identify major lineages. **(C)** Bar chart shows population fold-changes in relative abundance of each cluster. **(D)** UMAP as in Figure S5A split by sample. Color represents expression of Mki67. **(E)** Summary of cluster-by-cluster differential expression comparison of young and old samples. Each dot represents a gene, significant genes are shown in red. **(F)** Gene set enrichment analysis of significantly up- and down-regulated genes described in Figure S5E. **(G, H)** UMAP as in Figure S5A split by sample. Color shows average z-scores of genes in selected pathways.

**Figure S6. The lung RNA-sequencing analysis of old infected mice fed a chow or ketogenic diet, Related to Figure 6. (A)** Heatmap of normalized gene expression values of selected genes to identify major lineages. **(B)** UMAP plot of T cell cluster as in Figure 6C. Color represents expression of Trac. **(C)** Summary of cluster-by-cluster differential expression comparison of Old-KD and Old-Chow. Each dot represents a gene, significant genes are shown in red. **(D)** Percentage of each monocyte subset as identified in Figure 6F relative to total number of monocytes in the corresponding sample. **(E)** UMAP plot as in Figure 6F showing average z-scores of genes in IFN-alpha and IFN-gamma pathways.

